# *In vitro* ALPK1 kinase assay reveals new insights into ADP-heptose sensing pathway and kinase activity of disease-associated ALPK1 mutants

**DOI:** 10.1101/2023.01.04.522711

**Authors:** Diego García-Weber, Anne-Sophie Dangeard, Veronica Teixeira, Martina Hauke, Alexis Carreaux, Christine Josenhans, Cécile Arrieumerlou

## Abstract

Alpha-protein kinase 1 (ALPK1) is a pathogen recognition receptor that detects ADP-heptose (ADPH), a lipopolysaccharide biosynthesis intermediate, recently described as a pathogen-associated molecular pattern in Gram-negative bacteria. ADPH binding to ALPK1 activates its kinase domain and triggers TIFA phosphorylation on threonine 9. This leads to the assembly of large TIFA oligomers called TIFAsomes, activation of NF-κB and pro-inflammatory gene expression. Furthermore, mutations in *ALPK1* are associated with inflammatory syndromes and cancers. While this kinase is of growing medical interest, only few tools are available to characterize its activity. Here, we describe a versatile non-radioactive ALPK1 *in vitro* kinase assay based on protein thiophosphorylation. We confirm that ALPK1 phosphorylates TIFA T9 and show that T2, T12 and T19 are also weakly phosphorylated by ALPK1. Interestingly, we find that ALPK1 itself is phosphorylated in response to ADPH recognition during *Shigella flexneri* and *Helicobacter pylori* infection and that disease-associated ALPK1 mutants exhibit altered kinase activity. In particular, T237M and V1092A mutations associated with ROSAH syndrome and spiradenoma/spiradenocarcinoma respectively, exhibit enhanced ADPH-induced kinase activity and constitutive assembly of TIFAsomes. Altogether, this study provides new insights into the ADPH sensing pathway and a new tool for measuring the activity of ALPK1.

## Introduction

Alpha-protein kinase 1 (ALPK1, or lymphocyte alpha-kinase, LAK) is a kinase that is attracting growing interest because it recently appeared to be involved in important biological processes in health and disease. It belongs to the human alpha-kinase (α-kinase) family, a subset of six atypical kinases that share little sequence similarity in their catalytic domain with conventional kinases and which instead, have a kinase domain whose architecture is homologous to that of the myosin heavy chain kinases of *Dictyostelium discoideum* (Middelbeek et al. 2010). ALPK1 is a 139 kDa protein that contains an N-terminal alpha helical domain, an unstructured linker region and a C-terminal α-kinase domain. Like all members of the α-kinase family, it appeared recently in evolution and is only found in eukaryotes. ALPK1 was initially described as a protein involved in epithelial cell polarity and exocytic vesicular transport towards the apical plasma membrane (Heine et al. 2005). Indeed, Heine *et al*. showed that this kinase resides on golgi-derived vesicles where it phosphorylates myosin IA, an apical vesicle transport motor protein that regulates the delivery of vesicles to the plasma-membrane (Heine et al. 2005). ALPK1 has also been shown to phosphorylate myosin IIA and thereby modulate TNF-α trafficking in gout flares (Lee et al. 2016). More recently, our laboratory identified a role of ALPK1 in innate immunity during bacterial infection (Milivojevic et al. 2017). Indeed, we showed that this atypical kinase controls the oligomerization of TIFA and TRAF6 proteins, a process that activates the transcription factor NF-κB and induces the expression of inflammatory genes during infection by the Gram-negative bacteria *Shigella flexneri, Salmonella* Typhimurium and *Neisseria meningitidis* (Milivojevic et al. 2017). TIFA is a 20-kDa adaptor protein, which was first identified as a TRAF6-binding protein involved in NF-κB activation (Takatsuna et al. 2003). It contains a conserved threonine in position 9 (T9), a forkhead-associated (FHA) domain responsible for the recognition of phosphorylated threonine and serine residues and a glutamic acid at position 178 involved in TRAF6 binding. In resting conditions, TIFA is found as an intrinsic dimer (Huang et al. 2012). After phosphorylation by ALPK1 (Zhou et al. 2018), phosphorylated T9 (pT9) residues are recognized by FHA domains of adjacent dimers, leading to the formation of large TIFA oligomers, called TIFAsomes. We and others have shown that ADP-L-*glycero*-β-D-*manno*-heptose (ADPH), a soluble metabolite of the Gram-negative lipopolysaccharide (LPS) biosynthetic pathway, is the pathogen-associated molecular pattern (PAMP) whose recognition triggers the activation of the ALPK1/TIFA axis (Zhou et al. 2018; García-Weber et al. 2018; Pfannkuch et al. 2019). In addition, Zhou *et al*. showed that ALPK1 is the PRR for ADPH and characterized the binding site of ADPH on the N-terminal domain (NTD) of ALPK1 by crystal structure analysis (Zhou et al. 2018). They found that the NTD harbors 18 alpha-helices with 14 of them assembled into 7 antiparallel pairs that form a right-hand solenoid. ADPH binds in a pocket present on the concave side of the solenoid and several residues of ALPK1 interact with ADPH. Structural data suggest that the binding of ADPH to the NTD of ALPK1 interferes with the interactions that this domain has with residues of the kinase domain, which are predicted to be located near the kinase catalytic cleft (Zhou et al. 2018). This mechanism may induce a conformational change, which would make the catalytic cleft more exposed, thereby promoting the kinase activity of ALPK1. In addition to *S. flexneri, S*. Typhimurium and *N. meningitidis*, the ALPK1/TIFA axis is activated during infection with *Yersinia pseudotuberculosis* (Zhou et al. 2018), *Helicobacter pylori* (Stein et al. 2017; Gall et al. 2017; Zimmermann et al. 2017) and *Campylobacter jejuni* (Cui et al. 2021), showing that ALPK1 is involved in innate immunity against important human Gram-negative pathogens.

Beyond its role during infection, ALPK1 is also involved in the control of intestinal homeostasis. Indeed, ALPK1 was identified as the main intestinal inflammation regulator of the *Helicobacter hepaticus*-induced colitis and associated cancer susceptibility locus in mice (Ryzhakov et al. 2018). Ryzhakov *et al*. showed that in response to infection with the commensal pathobiont *Helicobacter hepaticus, Alpk1*-deficient mice display exacerbated interleukin-12 (IL-12)/interuleukin-23 (IL-23)-dependent colitis characterized by an enhanced Th1/interferon (IFN)-γ response. They showed that *Alpk1*, which is highly expressed by mononuclear phagocytes, controls intestinal immunity via the hematopoietic system. Indeed, in response to *H. hepaticus, Alpk1*^− /−^ macrophages produce abnormally high amounts of IL-12, but not IL-23 (Ryzhakov et al. 2018).

Finally, several publications reported a link between single-nucleotide polymorphisms or short deletion mutations in the *ALPK1* gene and disease susceptibility. Diseases include recurrent periodic fever (Sangiorgi et al. 2019), ROSAH syndrome (Williams et al. 2019), chronic kidney disease (Yamada et al. 2013), myocardial infarction (Fujimaki et al. 2014), ischemic stroke (Yamada et al. 2015), lung and colorectal cancer (Liao et al. 2016), spiradenoma and adenocarcinoma (Rashid et al. 2019), Type 2 Diabetes Mellitus and gout (Yamada et al. 2015; Ko et al. 2013) and interestingly, most of them share an inflammatory component. Whether mutations associated with these diseases alter the kinase activity of ALPK1 is unknown.

Although ALPK1 is involved in important biological processes, limited means are available to characterize its activity. To date, an antibody recognizing TIFA phosphorylated at its threonine 9 residue was the only tool available to monitor ALPK1 activity. However, since it is restricted to a specific phosphorylated residue, it does not allow the identification of additional TIFA phospho-sites or protein substrates. Here, we describe a versatile non-radioactive *in vitro* kinase assay based on protein thiophosphorylation reactions that allows the measurement of ALPK1 kinase activity towards any potential substrates. We confirmed that ALPK1 phosphorylates TIFA at position T9 and adressed the phosphorylation of additional TIFA residues. Interestingly, we discovered that ALPK1 itself is phosphorylated following ADPH recognition during pathogen exposure and that disease-associated ALPK1 mutants exhibit altered kinase activity.

## Results

### ADPH induces TIFA T9 phosphorylation, TIFAsome assembly and NF-κB activation in an ALPK1-dependent manner

In response to ADPH recognition, ALPK1 phosphorylates TIFA proteins on T9 residues (Zhou et al. 2018). Consecutively, inter-molecular binding of pT9 to FHA domains of TIFA dimers induces the formation of large TIFAsomes, a process that results in the oligomerization of TRAF6 and the activation of NF-κB. The formation of TIFAsomes can be visualized in HeLa cells expressing GFP-TIFA whereas the activation of NF-κB is measured by monitoring the nuclear translocation of p65 by immunofluorescence. The role of ALPK1 in ADPH sensing is confirmed in Figure 1 showing that the formation of TIFAsomes induced by ADPH treatment was inhibited in hALPK1 siRNA-transfected cells compared to control cells (Figure 1A and 1B). In line with this result, ADPH-induced NF-κB p65 nuclear translocation was also altered in ALPK1-depleted cells (Figure 1A) as quantified by measuring the cytosolic/nuclear p65 fluorescence intensity ratio by automated image analysis (Figure 1C). As expected, depleting ALPK1 did not affect TNFα-induced p65 nuclear translocation (Figure 1A and 1C). In order to directly address the role of ALPK1 in pT9 phosphorylation, HEK293 cells were transfected or not with a wild-type (wt) myc-hALPK1 cDNA construct and ALPK1 was immunoprecipitated using an anti-myc antibody. It was then incubated with purified GST-TIFA in the presence of ATP and analysed by western blot using the previously described anti-pT9 antibody (Zhou et al. 2018). Data confirmed that ALPK1 induced the phosphorylation of TIFA on T9 residues in response to the recognition of ADPH (Figure 1D and 1E). Altogether, these results confirmed that ADPH sensing induces TIFA T9 phosphorylation, TIFAsome assembly and subsequent NF-κB activation in an ALPK1-dependent manner.

**Figure 1:**
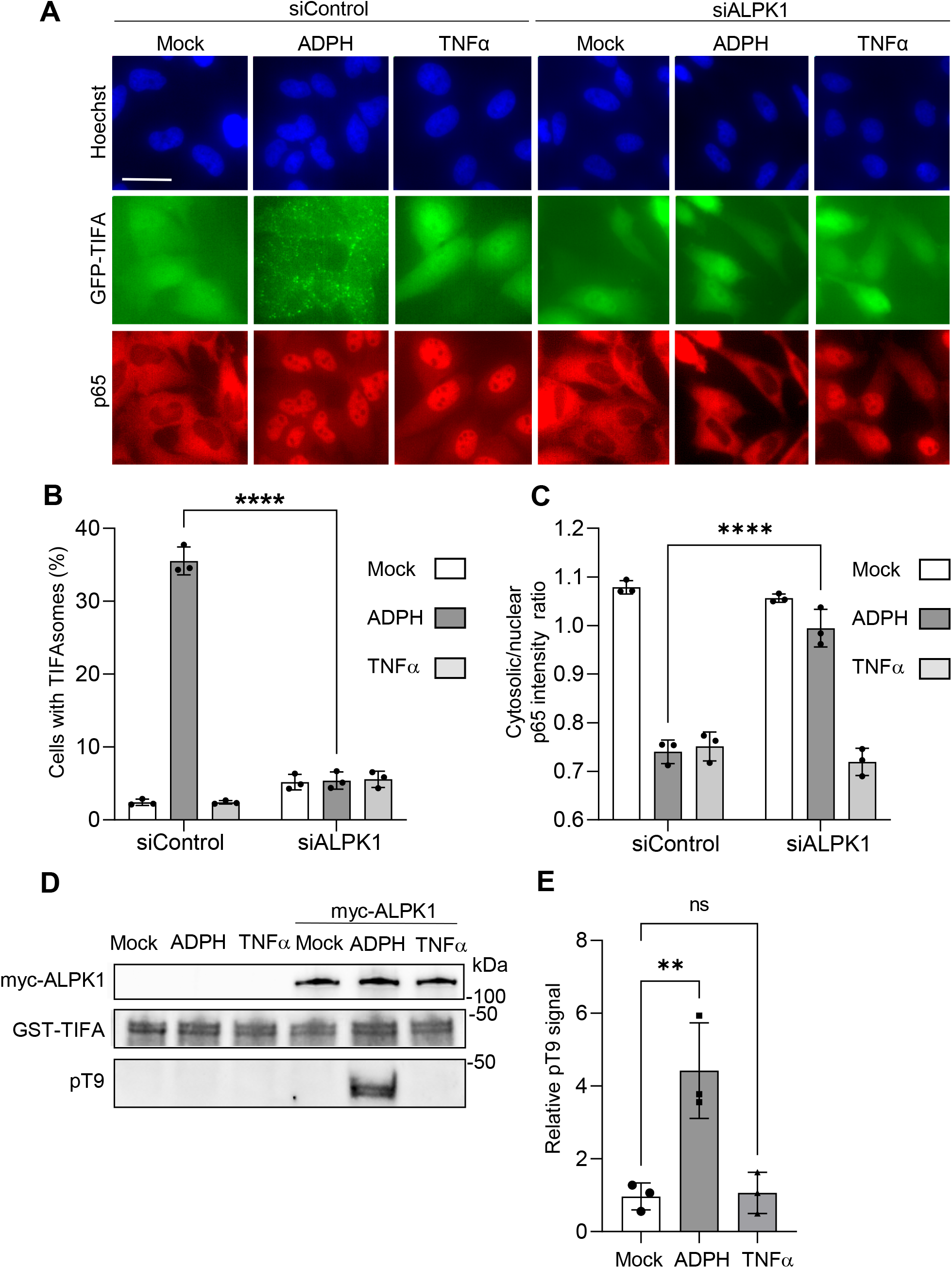
ADPH induces TIFA T9 phosphorylation, TIFAsome assembly and NF-κB activation in an ALPK1-dependent manner. **A**) ADPH-induced TIFAsome assembly and NF-κB nuclear translocation are ALPK1-dependent. Hela GFP-TIFA expressing cells were transfected with control or ALPK1 siRNAs and treated or not with ADPH or TNFα for 1 hour. They were then stained for p65 NF-κB (in red) and DNA with Hoechst (in blue). GFP-TIFA is shown in green. Bar scale is 10 μm. **B**) Quantification of the fraction of cells with TIFAsomes as shown in A. **C**) Quantification of p65 NF-κB nuclear translocation as shown in A. p65 translocation was quantified by measuring the p65 intensity ratio between the cytoplasm and the nucleus by automated image analysis. **D**) In response to ADPH sensing, ALPK1 phosphorylates TIFA at position T9. HEK293 cells were transfected or not with a wt myc-hALPK1 cDNA construct and stimulated or not with ADPH or TNFα for 1 hour. ALPK1 was immunoprecipitated and used in a GST-TIFA phosphorylation assay. myc-ALPK1, GST-TIFA and pT9 were detected by western blot with anti-myc, anti-GST and anti-pT9 antibodies, respectively. **E**) Quantification of TIFA T9 phosphorylation as shown in D. pT9 is quantified by measuring the ratio between pT9 and total GST-TIFA. All data correspond to the mean +/-SD of 3 independent experiments. Statistical significance was assessed using two-way ANOVA (B, C) followed by Tukey’s multiple comparisons test or one-way ANOVA followed by Dunnett’s multiple comparisons test (E) \*\**P* < 0.01, \*\*\*\**P* < 0.0001, not significant (ns).

### An *in vitro* kinase assay based on protein thiophosphorylation reveals the phosphorylation of ALPK1 in response to ADPH sensing

The development of an anti-pT9 antibody was decisive in the analysis of the ALPK1/TIFA axis. However, since its usage is limited to the analysis of TIFA phosphorylation on threonine 9 residues, it does not allow the identification of alternative phosphorylation sites on TIFA, nor any other ALPK1 potential substrates. For this reason, we developed an *in vitro* ALPK1 kinase assay, which uses ATPγS as phosphate donor and, which is based on the measurement of protein thiophosphorylation reactions. ATPγS is a non-hydrolysable ATP analogue that can be utilized by most kinases to transfer a thiophosphate group containing a sulfur atom at position gamma on a substrate. Briefly, HEK293 cells were transfected with a cDNA construct encoding myc-hALPK1 and, 48 hours later, were treated or not with ADPH. ALPK1 was immunoprecipitated with an anti-myc antibody and, to validate the assay, was incubated with its known substrate, purified GST-TIFA, in the presence of ATPγS at 37°C for 1 hour to allow protein thiophosphorylation. This critical step was followed by an alkylation reaction carried out by adding the alkylating reagent p-nitrobenzyl mesylate (PNBM) to the assay. Thus, thioester-type covalent bonds are formed at the locations occupied by the thiophosphate groups of the substrate and these can be detected by an anti-thiophosphate ester antibody (pTE) in western blots as previously described (Allen et al. 2007). As expected, ADPH treatment induced the thiophosphorylation of GST-TIFA and this process was dependent on the presence of ATPγS and the alkylating reagent PNBM in the assay (Figure 2A and 2B). It was also dependent on the presence of myc-ALPK1, strongly indicating that TIFA thiophosphorylation resulted from ALPK1 kinase activity (Figure 2C and 2D). Interestingly, a band corresponding to myc-hALPK1 was also detected by the anti-thiophosphate ester antibody, showing that ALPK1 was also thiophosphorylated in response to the recognition of ADPH (Figure 2A and 2C). As for TIFA, ALPK1 thiophosphorylation was dependent on ATPγS and PNBM. Interestingly, the mechanism of ALPK1 thiophosphorylation observed in response to ADPH sensing was independent on the presence of GST-TIFA in the assay (Figure 2C). In order to further characterize this new finding, we tested whether ALPK1 thiophosphorylation was dependent on the kinase activity of ALPK1. Cells were transfected with cDNA constructs encoding myc-ALPK1 wt, kinase (ΔK) or N-terminal (ΔN) domain deleted mutants, or with a K1067R mutant whose kinase acitivity is abolished (Zhou et al. 2018). As expected, ADPH-induced TIFA thiophosphorylation was abolished in cells expressing the different ALPK1 mutants (Figure 2E and 2F). Interestingly, ALPK1 thiophosphorylation was also abolished in these cells, showing that ADPH binding and ALPK1 kinase activity were both required for ALPK1 thiophosphorylation. Altogether, our ALPK1 activity assay based on protein thiophosphorylation confirmed that TIFA is a substrate of ALPK1, and revealed the phosphorylation of ALPK1 in response to ADPH sensing. Since the latter process is strictly dependent on the kinase activity of ALPK1, our results strongly suggest that the phosphorylation of ALPK1 results from an autophosphorylation mechanism.

**Figure 2:**
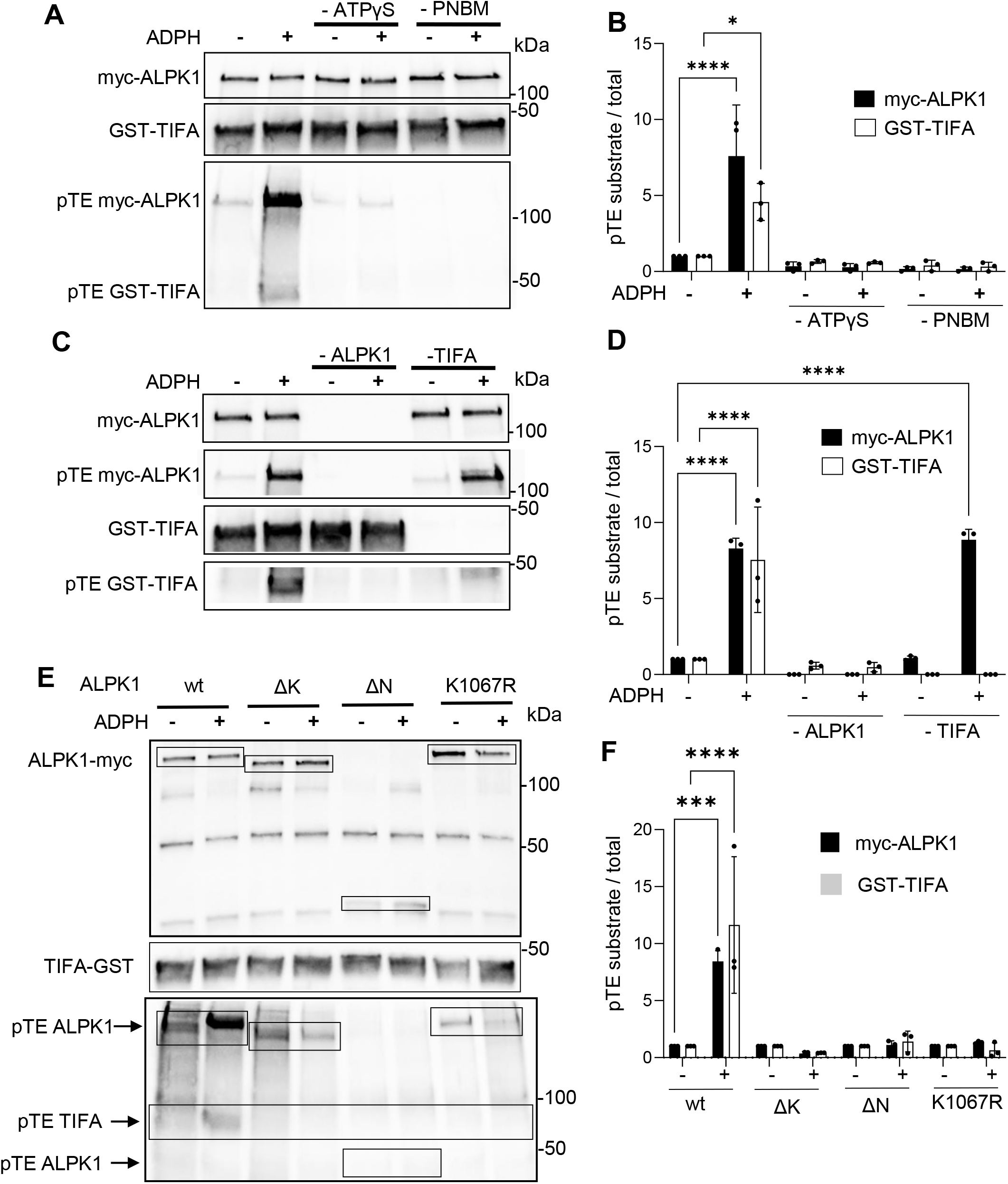
*In vitro* kinase assay based on protein thiophosphorylation reveals the phosphorylation of ALPK1 in response to ADPH sensing. **A**) TIFA and ALPK1 are thiophosphorylated in response to ADPH sensing. HEK293 cells, transfected with a myc-hALPK1 cDNA construct, were treated or not with ADPH for 30 minutes. myc-ALPK1 was immunoprecipitated and incubated with GST-TIFA +/-ATPγS and +/-PNBM. Anti-myc, anti-GST and anti-pTE antibodies were used in western blot to detect myc-ALPK1, GST-TIFA and thiophosphorylated proteins, respectively. **B**) Quantification of protein thiophosphorylation as shown in A. For GST-TIFA or myc-ALPK1, protein thiophosphorylation was quantified by measuring the ratio between corresponding pTE signal and total GST-TIFA or myc-hALPK1, respectively. **C**) Protein thiophosphorylation signals are dependent on ALPK1 and TIFA. HEK293 cells were transfected or not with a myc-hALPK1 cDNA construct and treated or not with ADPH. myc-ALPK1 was immunoprecipitated and incubated or not with GST-TIFA in the presence of ATPγS and PNBM. Anti-myc, anti-GST and anti-pTE antibodies were used in western blot to detect myc-ALPK1, GST-TIFA and thiophosphorylated proteins, respectively. **D**) Quantification of protein thiophosphorylation as shown in C. For GST-TIFA or myc-ALPK1, protein thiophosphorylation was quantified by measuring the ratio between corresponding pTE signal and total GST-TIFA or anti-myc, respectively. **E**) TIFA and ALPK1 thiophosphorylation is dependent on the kinase and N-terminal domains of ALPK1 and requires intact kinase activity. HEK293 cells were transfected with wt, ΔK, ΔN and K1067R myc-tagged ALPK1 cDNA constructs and were treated or not with ADPH. myc-ALPK1, GST-TIFA and thiophosphorylated proteins were detected as in A. **F**) Quantification of protein thiophosphorylation as shown in E. GST-TIFA and myc-ALPK1 thiophosphorylation was quantified as in B. All data correspond to the mean +/-SD of 3 independent experiments. Statistical significance against the untreated wt condition was assessed using two-way ANOVA (B, D, F) followed by Tukey’s multiple comparisons test \**P* < 0.05, \*\*\**P* < 0.001 \*\*\*\**P* < 0.0001.

### *S. flexneri* and *H. pylori* infections induce the phosphorylation of ALPK1

In order to test whether ALPK1 is phosphorylated during bacterial infection, HeLa cells were infected with the enteroinvasive bacterium *S. flexneri* at different multiplicities of infection (MOIs) and the thiophosphorylation status of GST-TIFA and myc-ALPK1 was analysed as described above. Consistent with data obtained in ADPH-treated cells, both proteins were increasingly thiophosphorylated with increasing MOIs (Figure 3A, 3B and 3C). Interestingly, their thiophosphorylation was sustained for several hours post infection (Figure 3D and 3E), indicating that the mechanism of ADPH sensing was likely active for this period in *S. flexneri* infected cells. TIFA and ALPK1 were also thiophosphorylated in epithelial cells infected with wild-type *Helicobacter pylori*, whereas an *hldE* gene deletion mutant (Δ*hldE*), unable to synthesize ADPH (Pfannkuch et al. 2019), failed to do so (Figure 3F, 3G and 3H). Altogether, these results showed that the mechanism of ADPH sensing that occurs during bacterial infection induces the phosphorylation of both TIFA and ALPK1.

**Figure 3:**
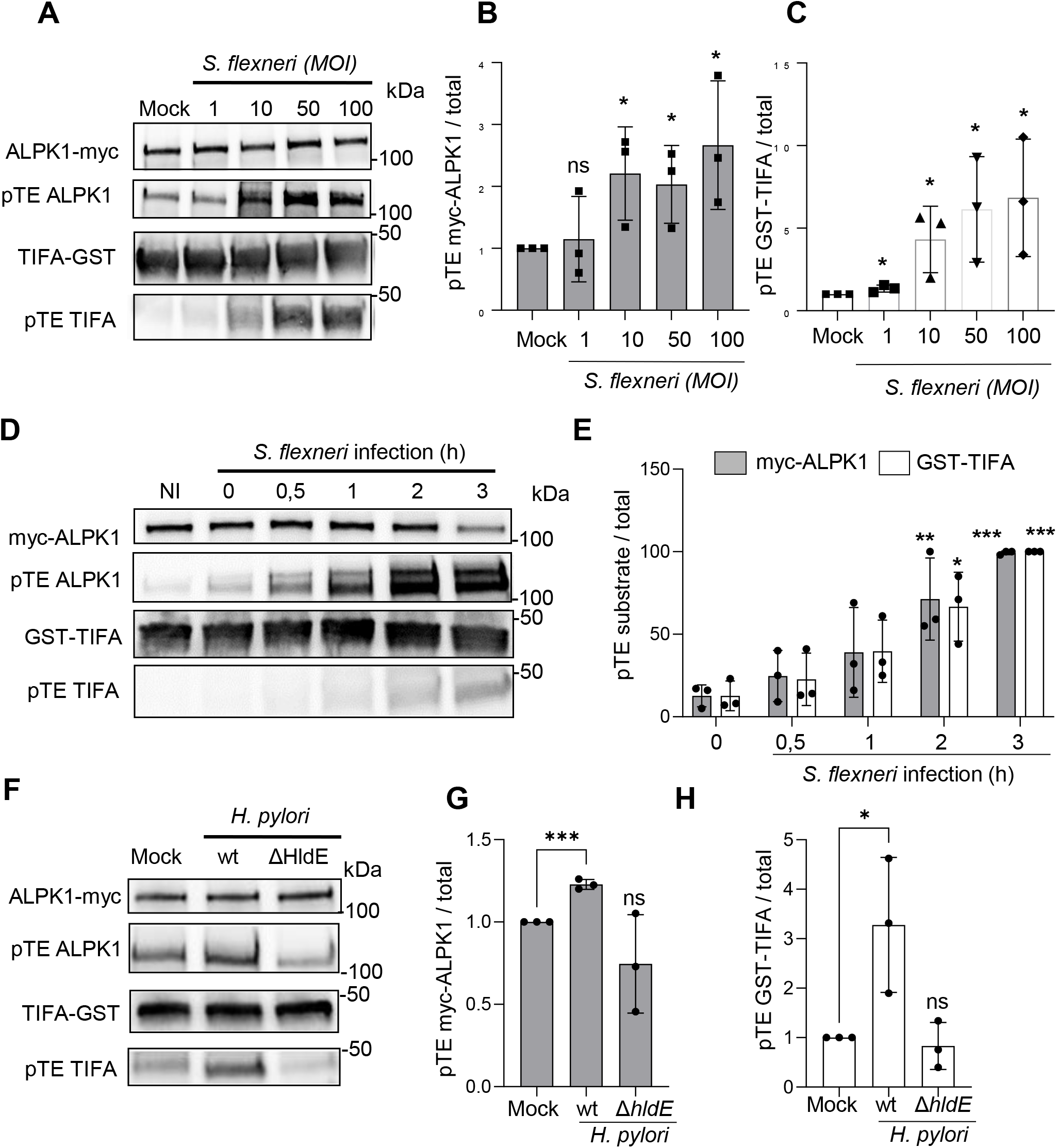
*S. flexneri* and *H. pylori* infections induce the phosphorylation of ALPK1 and TIFA. **A**) ALPK1 activity is enhanced in response to increasing *S. flexneri* MOIs. HEK293 cells, transfected with a myc-hALPK1 cDNA construct, were infected or not with wt *S. flexneri* at indicated MOIs for 1 hour. ALPK1 kinase activity was then assessed by the newly described protein thiophosphorylation assay. **B**) Quantification of myc-ALPK1 thiophosphorylation. **C**) Quantification of GST-TIFA thiophosphorylation. **D**) ALPK1 activity is sustainably enhanced during *S. flexneri* infection. HEK293 cells, transfected with a myc-hALPK1 cDNA construct, were infected or not with wt *S. flexneri* for indicated time-periods at MOI 10. ALPK1 kinase activity was then assessed by the newly described protein thiophosphorylation assay. **E**) Quantification of myc-ALPK1 and GST-TIFA thiophosphorylation. **F**) ALPK1 activity is increased during *H. pylori* infection of cells in an ADPH-dependent manner. HEK 293T cells transfected with a myc-hALPK1 cDNA construct, were infected or not with wt or an Δ*hldE H. pylori* mutant at MOI 20 for 4 hours. ALPK1 kinase activity was then assessed by the newly described protein thiophosphorylation assay. **G**) Quantification of myc-ALPK1 thiophosphorylation as shown in F. **H**) Quantification of GST-TIFA thiophosphorylation as shown in F. All data correspond to the mean +/-SD of 3 independent experiments. Statistical significance was assessed between mock and each condition individually using unpaired t-test (B, C, G, H) or using two-way ANOVA (E) followed by Tukey’s multiple comparisons test, \**P* < 0.05, \*\**P* < 0.01, \*\*\**P* < 0.001.

### ALPK1 phosphorylates TIFA on multiple residues

Since the newly developed assay allows to investigate ALPK1 kinase activity on any potential substrates, we further explored the substrate specificity of ALPK1. First, ALPK1’s ability to phosphorylate myelin basic protein (MBP) was monitored. MBP is a substrate commonly used in *in vitro* kinase assays, including those for MAPK, PKA, PKC cyclin-dependent and calmodulin-dependent protein kinases (Sanghera et al. 1990; Pietromonaco et al. 1998; Kishimoto et al. 1985). In contrast to TIFA, MBP was not thiophosphorylated in response to ADPH-dependent activation of ALPK1 (Figure 4A and 4B), suggesting that the kinase activity of ALPK1 is substrate specific. As TIFA is an important substrate of ALPK1, we also thoroughly explored its phosphorylation. In particular, we tested whether T9 is the only residue phosphorylated by ALPK1. To address this point, T9 and several amino acids surrounding T9 were individually mutated to alanine. Corresponding recombinant GST-TIFA proteins were produced in *Escherichia coli*, purified and used as substrates in thiophosphorylation-based ALPK1 kinase assay. As expected, thiophosphorylation was reduced for the T9A mutant, confirming that T9 is phosphorylated by ALPK1 (Figure 4C, 4D and 4E). Interestingly, it was not completely abolished, indicating that thiophosphorylation also occured on other TIFA residues. Quantification of normalized pTE GST-TIFA signals showed that thiophosphorylation was very slightly reduced for T2A, T12A and T19A but the effects were not statistically significant compared to wt (Figure 4E). To further characterize the potential phosphorylation of these residues, a T2A, T12A, T19A and T9A quadruple mutant was generated and used as ALPK1 substrate. When these four residues were mutated to alanine, thiophosphorylation was entirely abolished in a significant manner (Figure 4F and 4G), confirming that they were likely phosphorylated by ALPK1. In contrast to T9A, individual T2A, T12A and T19A mutants were all able to assemble into TIFAsomes in response to ADPH recognition during *S. flexneri* infection (Supplementary Figure S1). Taken together, these results show that, in addition to T9, several surrounding residues can be weakly phosphorylated by ALPK1. However, the phosphorylation of these residues is not necessary for the formation of TIFAsomes in response to ADPH sensing.

**Figure 4:**
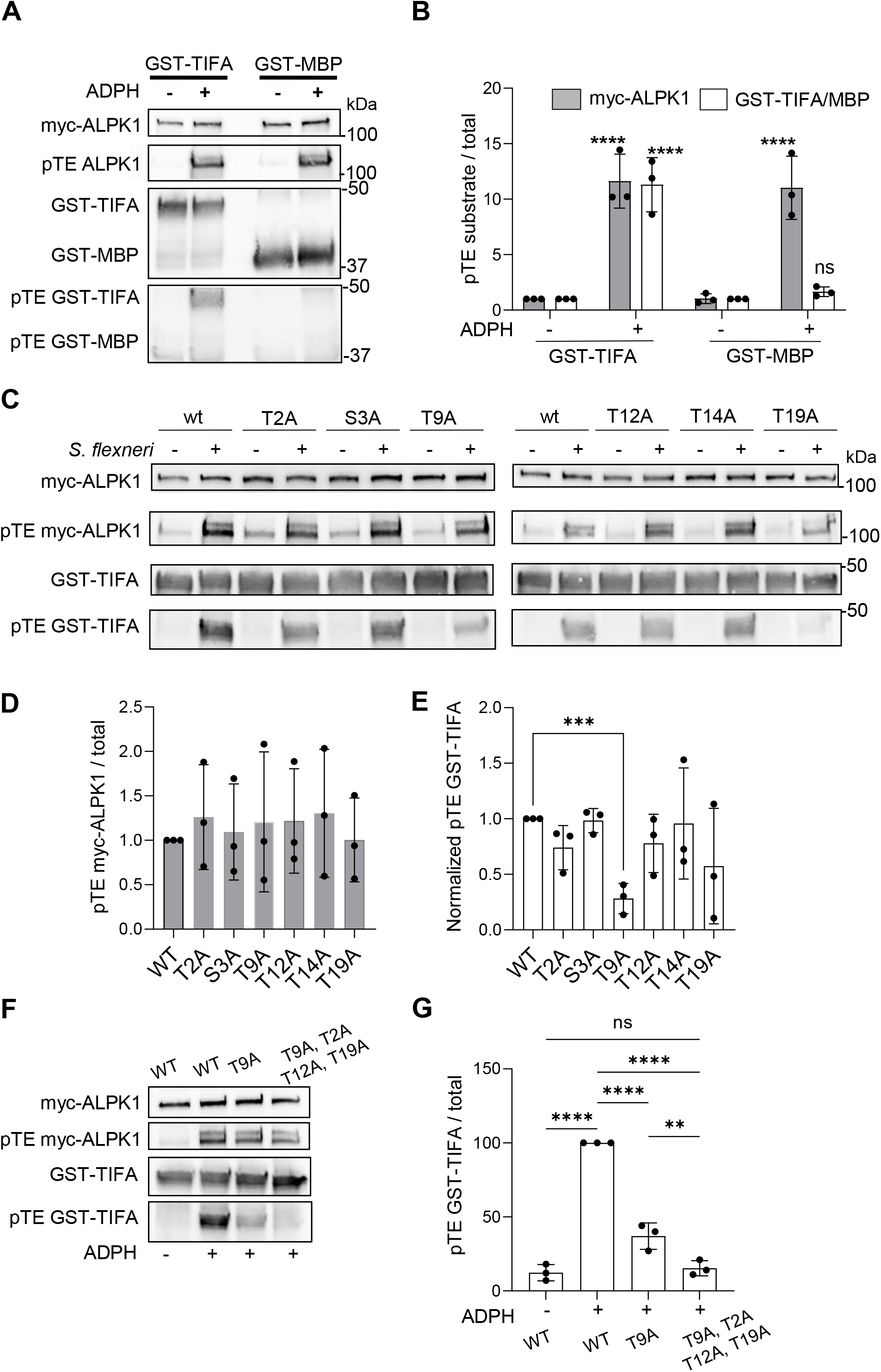
ALPK1 phosphorylates TIFA on multiple residues. **A**) MBP is not phosphorylated by ALPK1. HEK cells, transfected with a myc-hALPK1 cDNA construct, were treated or not with ADPH for 30 minutes. ALPK1 kinase activity was then assessed by incubating immunoprecipitated myc-hALPK1 with TIFA-GST or MBP. **B**) Quantification of myc-hALPK1, hTIFA-GST or MBP thiophosphorylation as shown in A. **C**) ALPK1 phosphorylates TIFA on multiple residues. HEK293 cells were transfected with a myc-hALPK1 cDNA construct, myc-hALPK1 was immunoprecipitated and incubated with wt, T2A, S3A, T9A, T12A, T14A or T19A GST-TIFA purified proteins. **D**) Quantification of myc-hALPK1 thiophosphorylation as shown in C. **E**). Quantification of GST-TIFA thiophosphorylation as showin in C. pTE GST-TIFA were normalised to corresponding total GST-TIFA and pTE myc-hALPK1 signals. Data correspond to the mean +/-SD of 3 independent experiments. **F**) ALPK1 phosphorylates TIFA on multiple residues. Cells were treated as in C. Thiophosphorylation assay was performed with wt, T9A and T9A/T12A/T14A/T19A GST-TIFA purified proteins as substrates. **G**) Quantification of GST-TIFA thiophosphorylation as showin in F. Data correspond to the mean +/-SD of 3 independent experiments. Statistical significance was assessed using one-way ANOVA followed by Dunnett’s multiple comparisons test (G), or two-way ANOVA followed by Tukey’s multiple comparisons test (D, E). Statistical significance was assessed between mock and each condition individually using unpaired t-test (D, E). \**P* < 0.05, \*\**P* < 0.01, \*\*\**P* < 0.001.

### Several disease-associated ALPK1 mutants have altered kinase activity

Mutations in ALPK1 have been recently described as associated or responsible for diseases, including inflammatory disorders and cancers. Therefore, we used the thiophosphorylation-based ALPK1 assay to test whether these mutations have an impact on the kinase activity of ALPK1 in resting conditions or after cell treatment with ADPH. For this, c.710C>T and c.3275T>C were introduced into a myc-hALPK1 Cdna construct by directed mutagenesis and corresponding p.T237M and p.V1092A ALPK1 mutants were generated. For each of them, the kinase activity was measured. An increase of thiophosphorylation of GST-TIFA and myc-ALPK1 was observed in response to ADPH sensing, indicating that these mutants exhibit enhanced ADPH-induced kinase activity. These last results are consistent with published studies showing that the T237M mutant has a pro-inflammatory phenotype (Williams et al. 2019) while V1092A activates NF-κB (Rashid et al. 2019). No significant increase in kinase activity was observed in absence of ADPH treatment (Figure 5A, 5B and 5C), suggesting that these mutants were not constitutively activated. However, since a bulk biochemistry assay may mask weak responses from some cells expressing ALPK1 mutants, we monitored ALPK1 activity with a single-cell functional assay. For this, the ability of T237M and V1092A ALPK1 mutants to trigger the constitutive formation of TIFAsomes was analysed. Hela cells stably expressing GFP-hTIFA were depleted of endogenous ALPK1 and transiently transfected with wt, T237M or V1092A myc-hALPK1 cDNA constructs. After fixation, cells were analysed by microscopy and the fraction of cells with TIFAsomes was quantified. Interestingly, some TIFAsomes were observed in a large fraction of cells expressing V1092A (Figure 5D and 5E). In addition, a few TIFAsomes were also visible in a small fraction of cells expressing the T237M mutant but this was not statistically significant (Figure 5D and 5E). Altogether, we showed that several ALPK1 mutants associated with diseases have an altered ADPH-induced kinase activity and that the V1092A mutant, more particularly associated with spiradenoma and spiradenocarcinoma, induces massive assembly of TIFAsomes in a constitutive manner.

**Figure 5:**
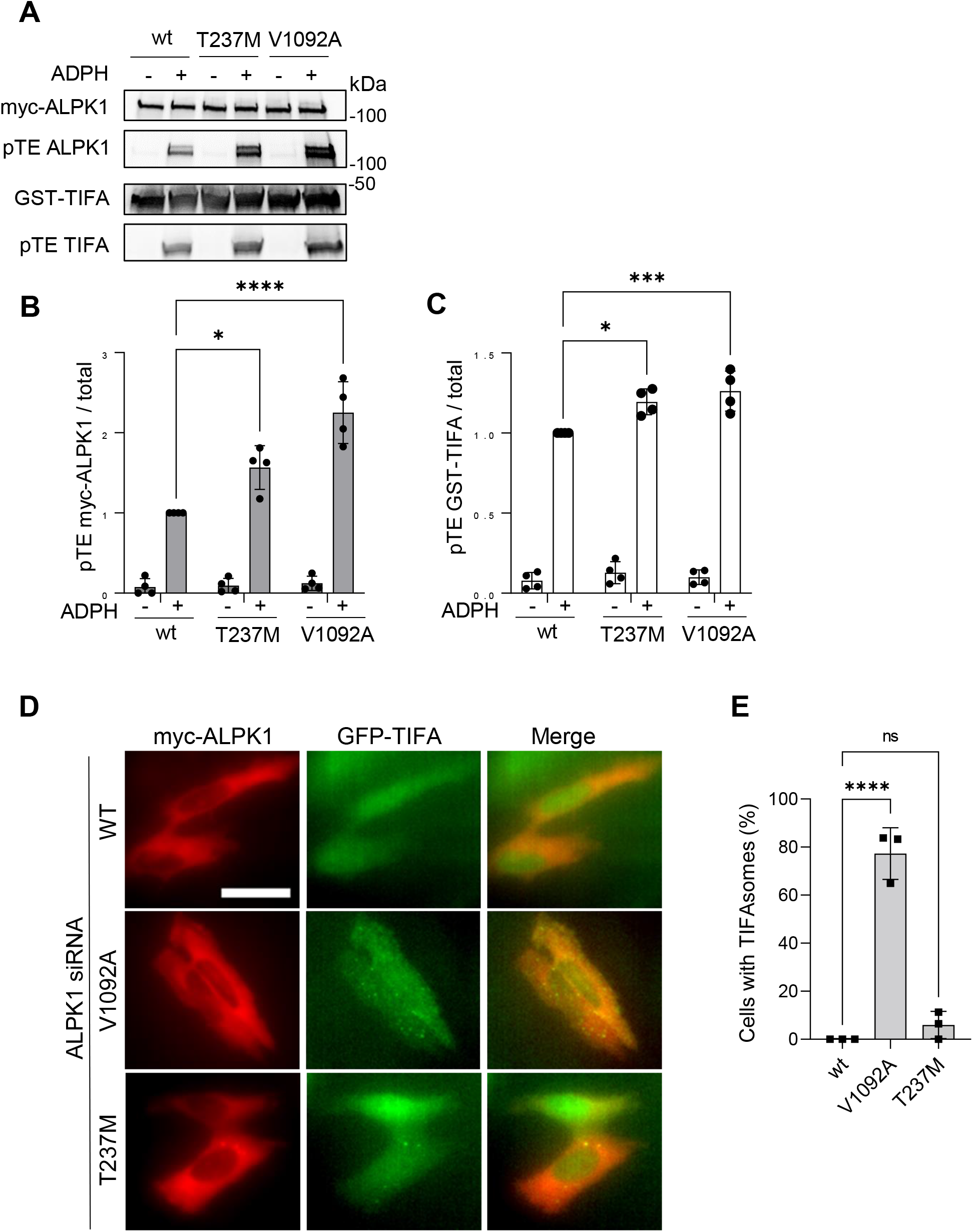
Disease-associated ALPK1 mutants have altered kinase activity. **A**) Disease-associated mutants have altered myc-ALPK1 and GST-TIFA thiophosphorylation. HEK293 cells transfected with wt or indicated mutated myc-hALPK1 constructs and treated or not with ADPH. ALPK1 kinase activity was then assessed using the newly described protein thiophosphorylation-based assay. **B**) Quantification of myc-hALPK1 thiophosphorylation as shown in A. **C**) Quantification of GST-TIFA thiophosphorylation as shown in A. **D**) V1092A mutant induces constitutive assembly of TIFAsomes. GFP-TIFA-expressing Hela cells were depleted of endogenous ALPK1 by siRNA and transfected with wt, T237M and V1092A myc-tagged hALPK1 constructs. After 12 hours of transfection, cells were fixed without ADPH treatment and stained for myc. TIFA is in green, myc-ALPK1 in red. Bar is 30 μm. **E**) Quantification of cells with TIFAsomes. For each construct, n>150 cells expressing a myc-hALPK1 construct were selected and the fraction of cells with TIFAsomes was quantified. Data correspond to the mean +/-SD of 4 (B, C) or 3 (D, E) independent experiments. Statistical significance was assessed using two-way ANOVA followed by Tukey’s multiple comparisons (B, C) or one-way ANOVA followed by Dunnett’s multiple comparisons test (E).

## Discussion

ALPK1 is a kinase that is receiving increasing attention in the field of biomedical research. Yet, very limited tools are available to study this protein. Here, we developed a versatile non-radioactive *in vitro* kinase assay that potentially allows to measure ALPK1 kinase activity towards any substrates whereas the currently existing method is limited to the analysis of the pT9 residue of TIFA. Our assay based on thiophosphorylation reactions confirmed that pT9 TIFA is a substrate of ALPK1. Interestingly, we found that ALPK1 itself is thiophosphorylated in response to ADPH treatment. Although we can not exclude that ALPK1 may be phosphorylated by another kinase that could be pulled-down with ALPK1 during the immunoprecipitation process, the observation that this mechanism depends on the kinase activity of ALPK1 strongly suggests that ALPK1 is autophosphorylated upon ADPH-induced activation. More work is needed to characterize the precise mechanism of autophosphorylation. In general, autophosphorylation can occur in cis when a kinase’s own active site catalyzes the phosphorylation reaction, or in trans when another kinase of the same kind provides the active site that carries out the enzymatic reaction. Determining whether ALPK1 can form dimers will be critical as trans-autophosphorylation is more frequent when kinase molecules dimerize. More work will also be needed to identify the residues that are autophosphorylated in response to ALPK1 activation. Analysis of ALPK1 activity during *S. flexneri* infection showed that this PRR is activated within minutes of infection and that activation is maintained for several hours, most likely reflecting sustained ADPH release from intracellular bacteria. This may result from residual lysis, active replication of intracellular bacteria or secretion via their T3SS, as previously suggested (Gaudet et al. 2017). The use of the ALPK1 *in vitro* kinase assay reveals that T9 is not the only residue that is phosphorylated by ALPK1 in response to ADPH sensing. Data showed that T2, T12 and T19 are also weakly phosphorylated by ALPK1. In response to TNFα stimulation, Huang et *al*. found that T9 is the only phosphorylated residue within the T9-containing peptide (^1^MTSFEDADTEE^11^), indicating that T2 is not phosphorylated (Huang et al. 2012). However, it should be noted that ALPK1 is not activated in response to TNFα (Zhou et al. 2018) and therefore, this is not the kinase that phosphorylates TIFA in this pathway. Interestingly, imaging experiments performed with alanine substitution mutants of T2, T12 and T19 residues revealed that they are not involved in the formation of TIFAsomes, confirming that T9 is the central residue in this important process observed in response to ADPH sensing. The impact of T2, T12 and T19 phosphorylation remains to be elucidated.

We used the newly developed kinase assay to measure the activity of disease-associated ALPK1 mutants. Interestingly, we found that both T237M and V1092A exhibit an increase of kinase activity with the latter having the strongest effect. In addition to this bulk assay, imaging experiments monitoring the formation of TIFAsomes in single cells in absence of ADPH treatment showed that V1092A, and in a much less extend T237M, can induce a constitutive formation of TIFAsomes. T237 is positioned within the ADPH binding site of ALPK1 wheras V1092 is located in the α-kinase domain. Results obtained with these mutants are in line with their phenotypes in the diseases to which they are associated. T237M is responsible for a NF-κB-mediated autoinflammatory disease in which nearly all patients exhibited at least one feature consistent with inflammation (Kozycki et al. 2022). V1092A, associated with spiradenomas, is also responsible for constitutive NF-κB activation (Rashid et al. 2019). Our data indicate that for both ALPK1 mutants, constitutive activation of NF-κB is likely due to constitutive formation of TIFAsomes.

In conclusion, this study provides a non-radioactive assay to analyse the kinase activity of ALPK1. It allows us to show that ALPK1 is likely autophosphorylated in response to ADPH recognition and that it can phosphorylate T9 but also additional residues of TIFA. In addition, this assay can be combined with an image-based single cell assay monitoring the formation of TIFAsomes to characterize the basal kinase activity of ALPK1 mutants associated with diseases. Since this assay is not focused to the phosphorylation of one protein in particular, it could be used to identify new substrates of ALPK1 and thereby characterize additional cellular pathways regulated by this kinase involved in health and disease.

## Methods

### Cell culture and reagents

HEK293 and HeLa cells (American Type Culture Collection) were cultured in Dulbecco’s modified Eagle’s medium (DMEM) supplemented with 10% fetal calf serum, 2 mM GlutaMAX-1, 100 μg/mL streptomycin and 100 U/mL penicillin (complete growth medium) at 37°C under 5% CO_2_. HeLa cells stably expressing GFP-TIFA were previously described (García-Weber et al. 2018). Cells were not cultured in the presence of antibiotics when co-incubated with live *H. pylori*. Pure ADP-heptose was purchased at J&K Scientific BVBA (#9020852) and Invivogen (tlrl-adph-I).

### Bacterial strains

DsRed-expressing wild-type (wt) M90T *S. flexneri* and wt and Δ*hldE H. pylori* strains were described previously (Milivojevic et al. 2017; Stein et al. 2017).

### ALPK1 and TIFA cDNA constructs

Wt or mutated human (h)ALPK1 cDNA constructs were cloned into the pCMV-myc plasmid (Takara Bio Inc) and were all siRNA resistant against the hALPK1 siRNA (s37074) from Ambion (Thermo Fisher Scientific) (Milivojevic et al. 2017). In the ΔN ALPK1 mutant (1017-1237), the N-terminal domain of ALPK1 was deleted. It was generated using SalI and NotI restriction enzyme sites which were added by PCR using primers listed in Table 1. The PCR product was then digested with Sal1 and Not1 and ligated into pCMV-myc vector. The ΔK ALPK1 mutant was also generated using SalI and NotI restriction enzymes for cloning. K1067R, T237M and V1092A mutants of ALPK1 were generated by using the QuikChange XL mutagenesis kit (Agilent Technologies, # 200516) and the primers described in Table 1. Wt and T9A GST-TIFA cDNA were generated by cloning into pGEX-4T3 plasmid (kindly provided by F.

**Table 1:**
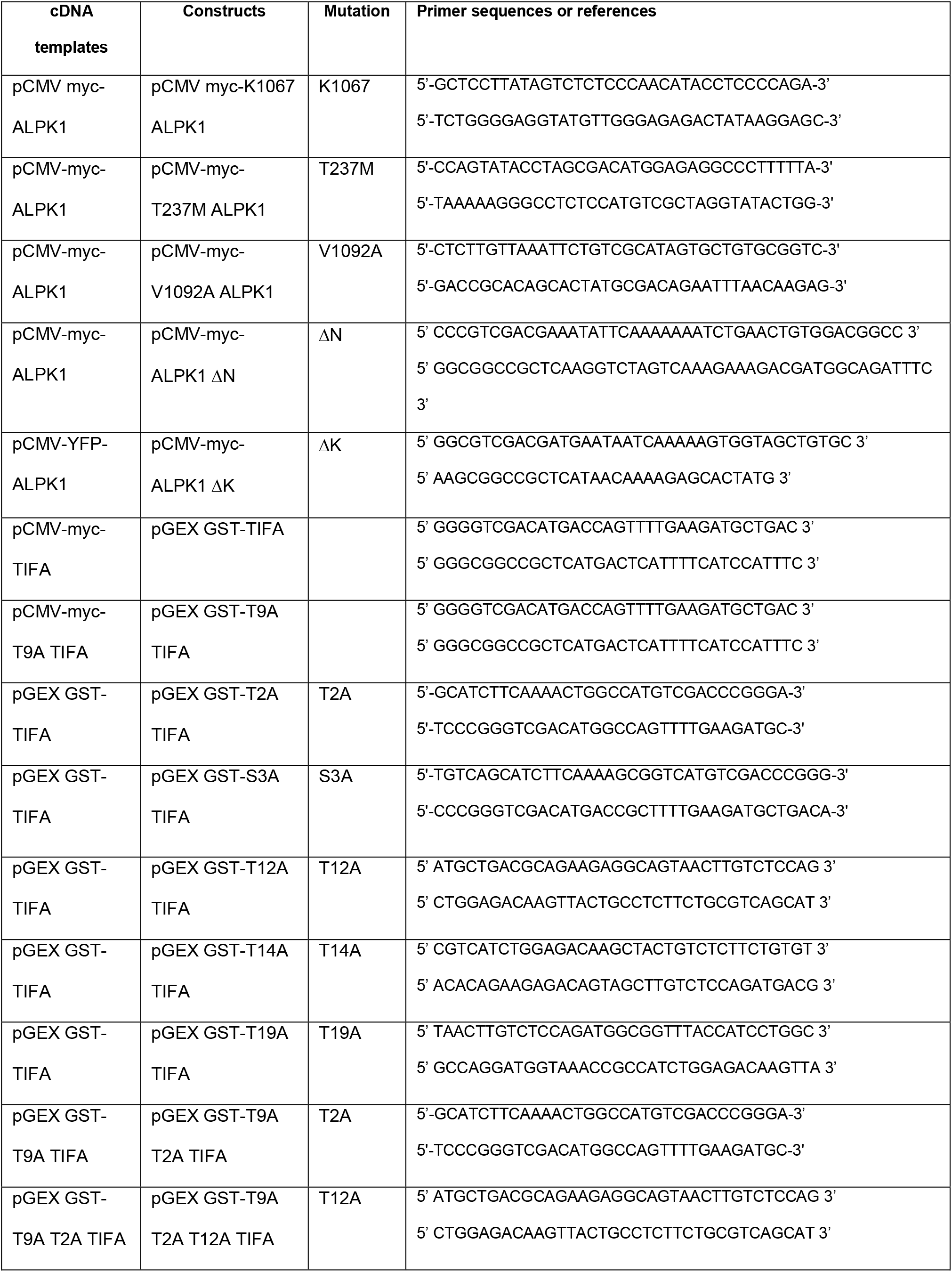
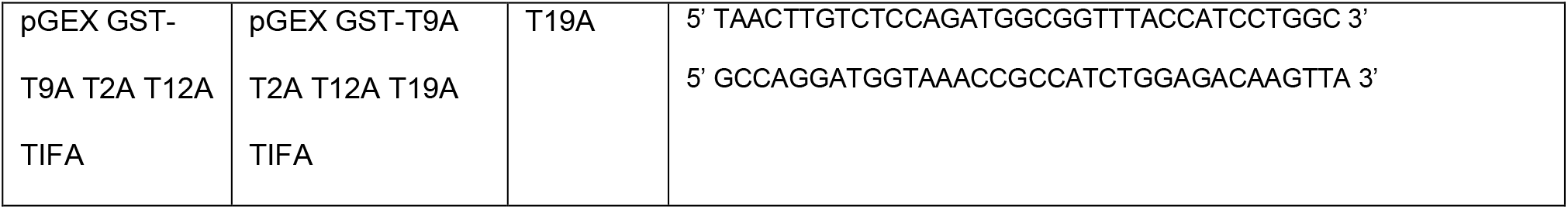
List of primers used to generate the ALPK1 and TIFA constructs.

Margottin, Institut Cochin, France). Wt and T9A pCMV-myc-TIFA cDNA constructs (Milivojevic et al. 2017) were used as PCR templates for cloning. Briefly, SalI and NotI restriction enzyme sites were added at each extremity of the genes by PCR using primers described in Table 1. PCR products were then digested with SalI and NotI and ligated into pGEX-4T3 vector. Single mutants T2A, S3A, T9A, T12A, T14A, T19A of TIFA were also generated with the QuikChange XL mutagenesis kit. Quadruple GST-TIFA T9A, T12A, T14A, T19A mutant was obtained by three sequential rounds of directed mutagenesis.

### Transfection and rescue experiments

HEK293 cells were seeded at 400,000 cells per well in 6-well plates and transfected the day after with indicated cDNA constructs with FuGENE6 (Roche). Reverse transfection of siRNAs was carried out using RNAiMAX according to the manufacturer’s instructions (Invitrogen Thermo Fisher Scientific). Briefly, HeLa cells, seeded in 96-well plates (8,000 cells/well), were reverse-transfected with 20 nM siRNA and used 72 hours post transfection. As previously described (Milivojevic et al. 2017), cells were transfected with a non-targeting siRNA sequence (4390843) or ALPK1 siRNA (s37074) from Ambion (Thermo Fisher Scientific). To replace endogenous ALPK1 by wt or mutated ALPK1, cells were first transfected with ALPK1 siRNA (s37074). Then, 48 hours later, they were transfected with siRNA-resistant wt, T237M or V1092A mutated hALPK1 cDNA constructs using FuGENE 6 (Roche).

### Purification of GST-TIFA proteins

Thermocompetent *E. coli* BL21 were transformed with wt and mutated pGEX-4T3-GST-TIFA constructs. For each construct, several colonies were cultivated in LB medium containing ampicillin at 100 µg/mL. When bacteria reached an OD_600_ value of 0.4, they were induced with 0.2 M IPTG and incubated at 16°C over night under agitation. Bacterial cultures were then centrifuged at 6 000 x g for 20 minutes at 4°C and pellets were frozen at -20°C overnight. They were then thawed and resuspended in cold lysis buffer containing PBS 1X, 2% Triton X-100, 1 mM PMSF and 2 mg/mL lysozyme (Sigma) and incubated for 1 hour on ice. Then, lysates were sonicated (3 pulses of 30 secondes with 30 second-breaks and 40% amplitude). Sonicated lysates were centrifuged at 16 000 x g for 30 minutes at 4°C. Supernatants were filtered using 0.45 µm syringue filter. Protein purification was performed using glutathione sepharose 4B resin according to the manufacturer’s instructions (GE Healthcare). Eluted proteins were then dialyzed and concentrated using an Amicon Ultra-15 10 KDa cutoff (Millipore).

### Permeabilized cell assay for ADPH treatment

Cells were treated as previously described (García-Weber et al. 2018). Briefly, cells were washed in permeabilization buffer containing 100 mM KCl, 3 mM MgCl_2_, 50 mM HEPES, 0.1 mM DTT, 85 mM sucrose, 0.2% BSA, and 0.1 mM ATP. They were then incubated with digitonin (2.5 μg mL_− 1_) plus ADPH at the desired concentration (J&K Scientific BVBA, #9020852) for 30 min in this same buffer. According to the experiment, cells were then fixed or washed and incubated in DMEM supplemented with 1% FCS for indicated durations.

### Bacterial infections

For *S. flexneri* infection, bacteria were used in exponential growth phase and coated with poly-L-lysin prior to infection, as previously described (Milivojevic et al. 2017). Cells were seeded in 6-well plates and infected at the indicated multiplicities of infection (MOIs) in DMEM supplemented with 10 mM HEPES, 2 mM GlutaMAX-1 and 1% FCS.

After adding bacteria, plates were centrifuged for 3 min at 300 x *g* and placed at 37°C for the indicated times. Cells were then lysed in 1 mL of 1% NP-40 lysis buffer and cell lysates processed for immunoprecipitation and *in vitro* kinase assay. *H. pylori* infection experiments were carried as previously described (Stein et al., 2017). Briefly, HEK 293 cells were seeded at 1 × 10^6^ cells per well in 6-well plates and let multiply over night to a confluency of the monolayer of roughly 50-70%. Cells were transfected with myc-hALPK1 construct or empty vector, using lipofectamine 2000 (Invitrogen) according to the manufacturer’s instructions. Cells were left to express the ALPK1 construct for 48 hours after transfection. Subsequently, freshly grown *H. pylori* (strain N6, wt and *hldE* mutant) was coincubated with the cells, in parallel with mock-infected controls, at an MOI of 20 bacteria per cell. After 4 h of coincubation, cells after medium removal were immediately harvested in 1% NP40-RIPA buffer (800 µl), lysed for 30 min on ice, snap-frozen in liquid nitrogen, and further processed for *in vitro* kinase assay and protein analysis. Cell protein content of the cleared lysates was determined using BCA assay. Uniform ALPK1 expression was subsequently monitored for all samples using Western immuno-blot on equal amounts of loaded samples with anti-hALPK1 or anti-myc antibodies. The experimental setup was repeated three times (biological replicates) on different days.

### *In vitro* kinase assay with ATPγS

HEK293 cells were seeded 72 hours prior to the experiment at 400,000 cells per well in 6-well plates and transfected the next day with wt or mutated myc-ALPK1 constructs with FuGENE6 (Roche) according to manufacturer’s instructions. The day of the experiment, cells were infected or stimulated with ADPH and cells were lysed in a lysis buffer containing 1% NP-40, 150 mM NaCl, 10 mM Tris, 5 mM EDTA, 10% Glycerol, 1 mM vanadate, and Complete Protease Inhibitors (Roche). Total lysates were kept on ice for 20 min and then centrifuged at 16,000 g for 30 min. Supernatants were incubated with 1 μg of mouse monoclonal anti-myc antibody (9E10, Santa Cruz Biotechnology) on a rotating wheel overnight at 4 ºC to immunoprecipitate myc-tagged ALPK1 proteins. The next day, lysates were incubated with Protein G Dynabeads (Invitrogen Thermo Fisher Scientific) previously equilibrated in lysis buffer on a rotating wheel at 4ºC for 1 hour. Immunoprecipitates were washed five times in lysis buffer with the magnet « Dynamag » (Invitrogen) and washed in 1 mL of kinase buffer containing 62.5 mM HEPES, 1.625 mM DTT, 46.9 mM MgCl_2_, 3.125 mM EGTA and 15.6 mM beta-glycero-phosphate. For a standard kinase assay reaction, the whole immunoprecipitate was resuspended in 60 μL of kinase buffer containing 1 μg of recombinant GST-TIFA and 0,15 mM ATPγS. This suspension was incubated for 30 min at 37 ºC on a rotating wheel. Then, EDTA (20 mM) and the alkylation reagent *para*-nitro-benzyl-mesylate (PNBM, Abcam) at a final concentration of 2,5 mM, were added. The suspension was incubated at room temperature on a rotating wheel for 1 hour and then 20 μL of Laemmli 4x containing 20 mM DTT and 2 mM vanadate were added. Samples were kept at -20ºC until immunoblot analysis was performed.

### Immunoblotting

Samples were boiled for 5 minutes at 95 ºC and subjected to SDS-PAGE. Immunoblotting was performed using primary antibodies diluted in phosphate buffered saline containing 0.1% Tween and 5% nonfat dry milk. myc-tagged ALPK1, GST-TIFA and thiophosphorylated proteins were detected with a mouse monoclonal anti-myc antibody (9E10, Santa Cruz Biotechnology), a rabbit polyclonal anti-GST antibody (Abcam, ab19256) and a rabbit polyclonal anti-thiophosphate ester antibody (Abcam, ab92570), respectively. HRP-conjugated secondary antibodies were purchased from GE Healthcare, Cell Signaling technologies or ThermoFisher Scientific. Blots were analysed with an enhanced chemiluminescence method (SuperSignal West Pico Chemiluminescent substrate, Thermo Fisher Scientific).

### Immunofluorescence

After fixation in 4% paraformaldehyde, GFP-TIFA-expressing cells were incubated overnight at 4°C in 0.2% saponin with a mouse monoclonal anti-myc antibody (9E10, Santa Cruz Biotechnology) or with a mouse monoclonal NF-κB p65 antibody (Sc-8008). The day after, cells were washed five times with PBS 1X and incubated with secondary antibodies and Hoechst (Life Technologies, H3570) diluted in 0.2 % saponin for 45 min. Cells were washed five times in PBS 1X and analyzed by microscopy.

### Automated microscopy and image analysis

Images were acquired with an ImageXpress Micro (Molecular Devices, Sunnyvale, USA). Quantification of p65 nuclear translocation and cell fraction with TIFAsomes was performed using MetaXpress as previously described (Milivojevic et al. 2017; García-Weber et al. 2018). Each data point corresponds to triplicate wells, and more than nine images were taken per well.

### Statistical analysis

Data are expressed as mean ± standard deviation of at least three biological replicates. Tests used to assess statistical significance are described for each figure.

## Acknowledgements

We gratefully acknowledge financial support from the Agence Nationale pour la Recherche (grants no ANR-14-ACHN-0029-01 and ANR-17-CE15-0006 including postdoctoral fellowships to DGW) and from Fondation ARC pour la Recherche sur le Cancer (grant no ARC—PJA20171206187). We also acknowledge financial support through grant SFB 900 (project no: 158989968), sub-project B6 to CJ by the German Research Foundation. MH was additionally supported by the intramural graduate program “Infection Research on Human Pathogens@MvPI” established at LMU Max von Pettenkofer Institute. We thank Bettina Sedlmaier for expert technical assistance.

## Author contributions

DGW, CJ and CA designed research. DGW, ASD, VT, MH and AC performed research and analyzed data. DGW and CA wrote the manuscript. All authors discussed the results, edited and commented on the article.

## Supplementary data

**Supplementary Figure S1:**
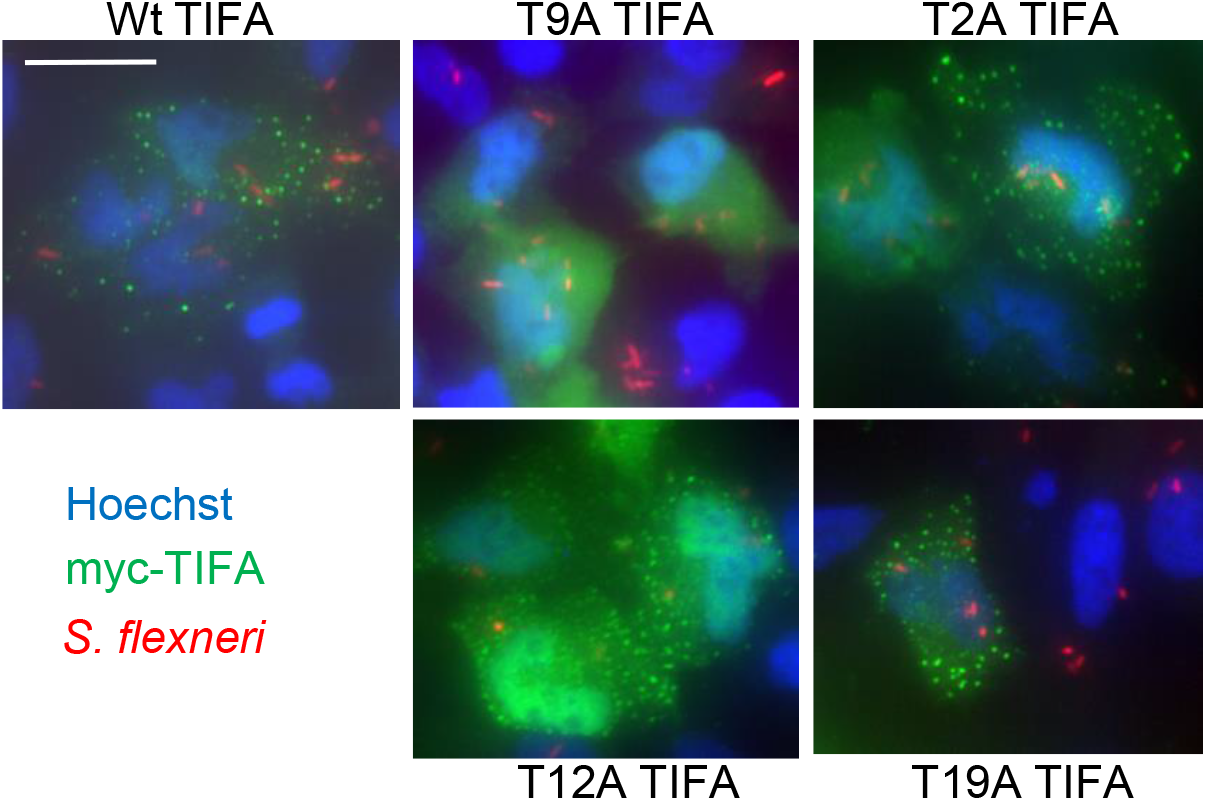
T2A, T12A and T19A TIFA mutants are able to assemble into TIFAsomes during *S. flexneri* infection. Hela cells were transfected with wt, T9A, T2A, T12A and T19A myc-TIFA constructs and infected with dsRed-expressing *S. flexneri* at MOI 10 for 1 hour. After fixation, cells were stained for myc (shown in green) and with Hoechst (shown in blue). Bacteria are shown in red. Bar is 30 μm.

